# Population estimation of a cryptic moss frog using acoustic spatially explicit capture recapture

**DOI:** 10.1101/2022.04.01.486679

**Authors:** Debra Stark, Andrew Turner, Berndt J. van Rensburg, John Measey

**Affiliations:** School of Biological Sciences, Centre for Biodiversity and Conservation Science, The University of Queensland, St Lucia, Queensland 4072, Australia; Centre for Invasion Biology, Department of Botany & Zoology, Stellenbosch University, Stellenbosch, South Africa; CapeNature Scientific Services, Private Bag X5014, Stellenbosch, 7599, South Africa; Biodiversity and Conservation Biology, University of the Western Cape, Private Bag X17, Bellville, 7535, South Africa; Department of Zoology, University of Johannesburg, Auckland Park, Johannesburg, 2006, South Africa

**Keywords:** Alien invasive plants, acoustics, off-reserve matrix, fire, spatial capture recapture, population estimation

## Abstract

Cryptic amphibians pose a problem for conservation managers as they are difficult to find to assess initial populations, and monitor changes during potentially threatening processes. The rough moss frog, *Arthroleptella rugosa*, is small and occurs in seepages on a single unprotected mountain in South Africa’s fire prone, biodiverse fynbos biome. The area is heavily impacted by invasive plants, which dry seepages and increase the frequency and intensity of fires, leading to the assessment of this species as Critically Endangered. We aimed to test the efficacy of acoustic spatially explicit capture recapture (aSCR) to estimate the entire population of calling adult *A. rugosa*, and assess the impacts by invasive plants. Using aSCR, our estimates suggest that the population of *A. rugosa* is more than five times that previously estimated using aural calling surveys on the mountain, at ∼2000 individuals. This despite an intense fire over the entire area three years earlier that reduced the calling population to a few tens of individuals. Our vegetation surveys suggest that the ongoing removal of invasive plants from the mountain is successful in areas occupied by *A. rugosa*, but that adjacent areas invaded by pines and hakea have a negative impact on calling density. The private public conservancy partnership on Klein Swartberg Mountain, is conserving this frog but will require ongoing management and monitoring to ensure conservation in the future.

## INTRODUCTION

Globally, threatened and range restricted species are of high conservation concern (IUCN 2018a; Gaston & Fuller 2009). Species with a restricted geographic range are often habitat specialists and weak dispersers, and thus face an increased risk of extinction due to threatening processes such as climate change, human development, and the spread of invasive species and diseases (e.g. Amphibians - Sodhi et al 2008; Cooper et al 2008; Mammals - Cardillo et al 2008; Birds, Lee & Jetz 2011; Plants - Casazza et al 2014). The rapid and increasing changes to habitat suitability and fragmentation are likely to, and in some cases already have, exceeded restricted-range species migration capabilities (Pearson 2006; Casazza et al 2014). In some countries, species listed as threatened by the IUCN (International Union for Conservation of Nature) are often afforded protection that requires impact mitigation through the protection, restoration and/or creation of habitat (Rodriguez *et al.* 2006), but in others no degree of threatened status affords any tangible protection level. Despite a global bias towards protecting range-restricted and threatened species, there is still a tendency for threatened species to be poorly represented in protected area networks across the globe (Rodrigues *et al.* 2004; Nori *et al.* 2015). Amphibians in particular, are the least represented taxon, with almost a quarter (1,535 species) of all known extant species (n. 6,500) remaining unrepresented in protected areas (Venter *et al.* 2014; Nori *et al.* 2015; IUCN 2018b), and most of these unrepresented species occur in only one site (Ricketts *et al.* 2005; Global Amphibian Assessment). Adding to the complexity of amphibian conservation is our general lack of understanding of the conservation status of those species or communities typically found within the off-reserve matrix (i.e., those typically excluded from reserve networks). Funding in the off-reserve matrix is often far more limited compared to resources linked to specific protected area management activities. Furthermore, monitoring rare and range-restricted species that require large sampling efforts can be difficult and impractical.

In South Africa, the most threatened native amphibian species are concentrated within the Cape Floristic Region, many of which are affected by agriculture (50%), invasive species (37.1%) and habitat change and loss (25.9%) (Stuart *et al.* 2004; Mokhatla *et al.* 2012; Angulo *et al.* 2011). Within this 8.77 million hectare (ha) region, protected areas cover over 2.3 million ha (26.6%) of land, and 96,557 ha (1.1%) of land comprised of privately-owned stewardship areas (Rouget *et al.* 2014). In these privately-owned areas there is a recognized need to integrate monitoring for improved conservation in an adaptive management framework (Rouget *et al.* 2014). Further to this, over half of the seepage habitat that many amphibian species rely on are Critically Endangered, with only 10% considered to be well protected (Driver *et al.* 2012). Annually, invasive plant species alone cause a loss of up to 87 million m^-3^.yr^-1^ of mean annual surface water runoff across South Africa (Wilson *et al.* 2014), which can be a potential fatal change for native amphibians. With only 12.5% of the natural threatened habitat area remaining in the Western Cape, the control of invasive species has been prioritized and is estimated to exceed an annual cost of US$ 23.5 million (Wilson *et al.* 2014).

Many of the amphibian species within South Africa are poorly studied, and their population responses to these threats remain poorly understood (Measey 2011; Measey *et al.* 2019). *Arthroleptella rugosa*, a Critically Endangered moss frog (IUCN & SA-FRoG 2017), is restricted to a single mountain in the off-reserve matrix of the Western Cape. It is currently threatened by invasive plants (*Pinus pinaster* and *Hakea sericea*), which degrade and dry their seepage habitat, and by fires in the invaded fuel-laden vegetation that are more severe than those it evolved in (Turner & Channing 2008; IUCN & SA-FRoG 2017). The synergistic interaction between these threats is thought to severely impact *A. rugosa* populations. As *A. rugosa* occurs exclusively on private land, it is dependent on the continued conservation efforts of the ‘Klein Swartberg Conservancy’; a collective of private landowners who designated the entire 14,857 ha mountain as a conservation area. CapeNature, the provincial nature authority, undertakes annual monitoring on the mountain to assess specific populations within the range of *A. rugosa*. Using aural calling surveys of the mountain (Dorcas *et al.* 2009), these survey techniques estimated the known populations to be around 400 adults (Turner & Channing 2008). This figure was doubled to account for females and rounded up to account for males that would not have been encountered during the brief survey to derive the estimate for the species of 1 000 individuals in the IUCN Red List assessment (IUCN & SA-FRoG 2017). An intense fire over the entire mountain in January 2012 was followed by further aural acoustic surveys that suggested the population was heavily impacted, with only a few tens of individuals calling.

Conventional amphibian population monitoring is typically carried out through visual surveys, trapping methods (capture-mark-recapture; CMR), or through acoustic surveys of calling males (Driscoll 1998; Dorcas *et al*. 2009; Marsh *et al*. 2017). Although these methods are commonly used and often largely successful, some can be impractical for cryptic or small species, particularly for those with small population sizes, such as *A. rugosa*. An acoustic approach, vocally ‘capturing’ individuals, is less invasive and can be more suited to these hard-to-find species. Calling surveys, completed manually by trained observers (such as those already conducted for *A. rugosa*), or through the use of automated recording systems provide rapid data that can be used to identify unnamed species, map distributions for occupancy models and deduce qualitative count data (Dorcas *et al*. 2009; Marsh *et al*. 2017). Aural calling surveys, however, can be subject to imperfect detection, misidentification and significant observer inconsistencies if conducted by inadequately trained observers (Dorcas *et al*. 2009). Observation errors such as omission (failure to detect a species that is present) and commission (incorrectly ‘detecting’ an absent species) (Parris *et al*. 1999; Tyre *et al*. 2003) make it difficult to reliably measure species occurrence or population size, which can significantly impact population models and derived population estimates (Rogers *et al*. 2013). The use of recording systems eliminates the dependence on real-time processing by skilled observers and can be used to collect permanent records of data, which can be examined and verified later (Dorcas *et al*. 2009). The applications of estimates attained using both aural calling surveys and traditional recording systems, however, are limited as the sampling area cannot be clearly defined and thus no density or actual population size estimates can be deduced; only qualitative animal counts within an unknown area (Stevens *et al*. 2002; De Solla *et al*. 2006).

Spatial capture-recapture (SCR, Efford 2004; Borchers 2012; Borchers & Fewster 2016) is a more recently developed method that combines capture-mark-recapture (CMR) and distance sampling methods (Buckland *et al.* 2001; Stevenson *et al.* 2015). While spatial capture-recapture was originally designed as a physical trapping method, it has since been adapted to use acoustic techniques to efficiently and non-invasively collect and analyze large volumes of acoustic data (Dawson & Efford 2009; Efford *et al.* 2009; Marques *et al.* 2012; Stevenson *et al.* 2015; Measey *et al.* 2017). This adapted technique, ‘acoustic spatial capture-recapture’ (aSCR), uses an array of fixed microphones to estimate the population density of vocalizing individuals and could be a suitable replacement for traditional survey methods typically used for visually cryptic, threatened and vocally distinct anuran species (Efford *et al.* 2009; Stevenson *et al.* 2015). While aSCR relies solely on calling individuals to determine population densities, these density estimates have been shown to be comparable to CMR estimates and thus can be reliable indicators of population trends and sizes (e.g. Meuche & Grafe 2005). Currently, aSCR is the only known acoustic density estimation method that can also generate confidence intervals in a statistically rigorous manner (Measey *et al.* 2017).

We implement this non-intrusive aSCR technique to accurately and efficiently estimate (i) the total adult male *A. rugosa* population across their range and (ii) *A. rugosa* adult male population densities in sites currently monitored and managed by conservation managers and private land owners. We assess the recovery of the population after the intense fire in 2012, and we use vegetation surveys in and adjacent to our recording areas to assess the efficacy of ongoing conservation efforts to control invasive plant populations. Population data will be used to assess the current *A. rugosa* population and conservation status, and direct ongoing and future efforts of private landowners and conservation managers towards areas of high importance for this species.

## MATERIALS AND METHODS

### Study Species

*Arthroleptella* is a genus of moss frogs within the speciose family Pyxicephalidae that are endemic to sub-Saharan Africa (Van der Meijden *et al.* 2011; Rebelo & Measey 2019). *Arthroleptella rugosa* is the most threatened and range-restricted species within this genus, occurring exclusively on private land on the Klein Swartberg Mountain (see Fig 1). *Arthroleptella rugosa* are typically found within, or in close proximity to, seepages or wetland flats where the soil is moist and there is adequate vegetation to provide protection from strong winds and high temperatures. Adults are small (mean body length of 13 mm; Turner & Channing 2008) with a dark brown appearance. Their size and cryptic coloration makes finding individuals difficult, however, like other *Arthroleptella* species; *A. rugosa* males are easily distinguishable and detectable by their advertisement calls (Turner & Channing 2017).

**Figure 1.**
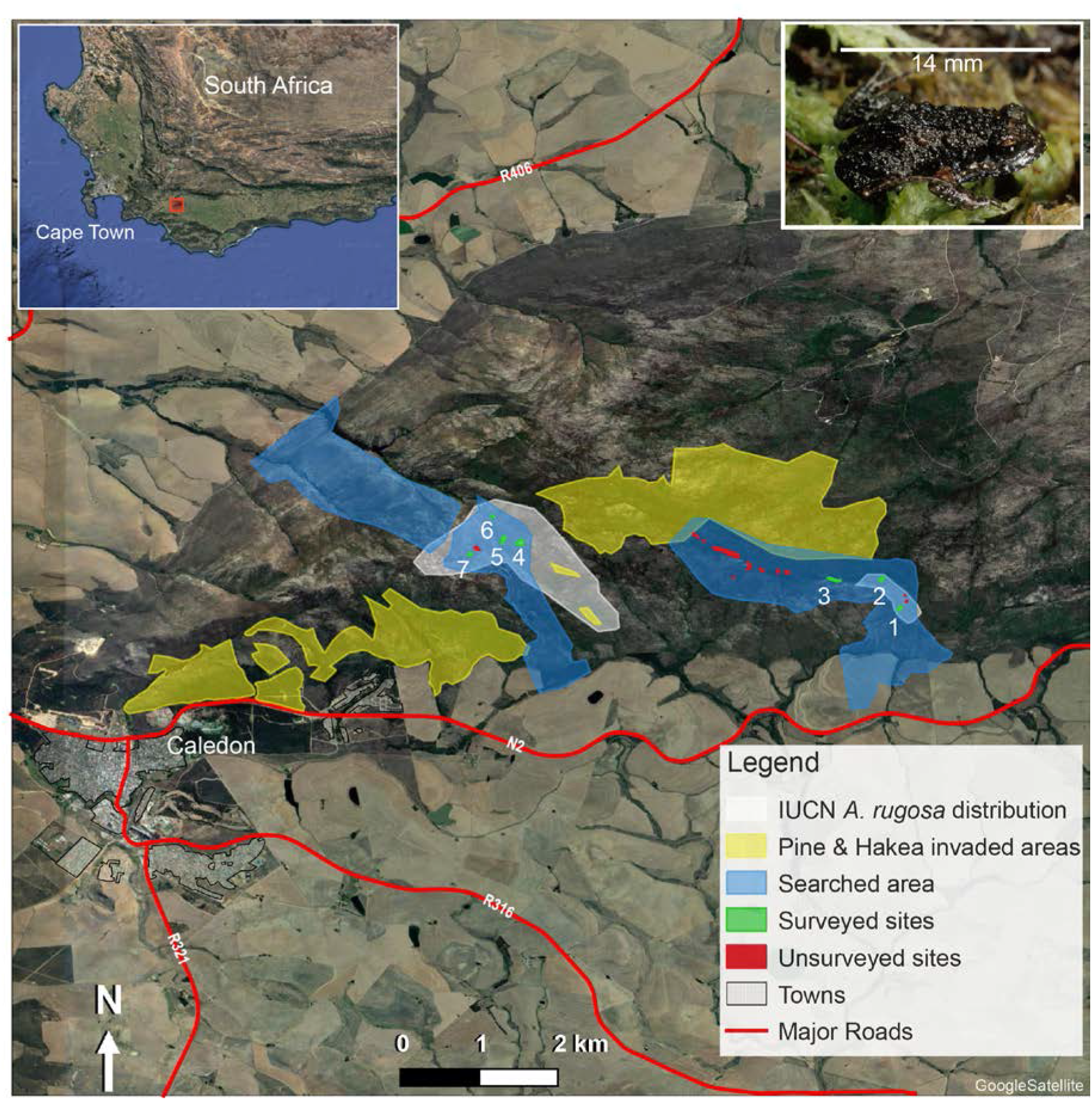
Map of the study area, with inset (top left) showing the position of the site in the southwest of South Africa, and (top right) a male *Arthroleptella rugosa* male with SVL 14 mm. The map demonstrates the search effort (blue polygons), identified areas of *Pinus pinaster* and *Hakea sericea* invasion (yellow polygons), surveyed habitat patches (green polygons), unsurveyed habitat patches (red polygons), and the IUCN listed *A. rugosa* habitat (grey polygons). Survey areas 1-3 are within the Eastern portion of the distribution and survey areas 4-7 are located within the Western portion.

*Arthroleptella* males are known to call from within seepage areas during an extended period throughout the austral winter; typically, May to November (Measey *et al.* 2017). *Arthroleptella rugosa* males exhibit three distinct vocalizations: a chirp-like advertisement call, a low frequency aggressive call and a chuckling call unique to this species (Turner & Channing 2008). *Arthroleptella rugosa* calling behavior is largely unknown, however, during the breeding season, males are known to frequently vocalize advertisement calls, as these are presumed to be used for attracting female mating partners and delineating territorial boundaries to conspecifics (Turner & Channing 2008). Advertisement calls have been shown to be the most frequently vocalized calls and are typically easy to capture in recordings (Kohler *et al.* 2017).

The entire population of *A. rugosa* occurs within a 230 ha area, and has previously been estimated as 400 adults, although this is expected to be in decline due to the ongoing plant invasion of *Pinus pinaster* and *Hakea sericea* (Turner & Channing 2008; SA-FRoG & IUCN 2017). A fire in January 2012 burnt the entire mountain (Measey *et al.* 2019; Turner 2012), with the following aural acoustic survey suggesting the *A. rugosa* population was heavily impacted with only a few tens of individuals calling in two areas of the eastern portion of the distribution (17 May 2012; AAT & JM pers. obs.). The increasing presence of alien plants reduces the availability of water, and increases the frequency and intensity of fires that naturally cycle in this area (le Maitre *et al.* 2002; Measey 2011). Consequently, a focused effort to remove alien vegetation from the mountain was started in 2012 under the auspices of CapeNature and supported by the US Fish and Wildlife Service. Due to the level of threat currently facing this species, its conservation status and the limited scientific literature available on the ecology of this species, *A. rugosa* has been identified as a high priority species requiring further research (Measey 2011).

### Study Area

The study area was located on private lands on the Klein Swartberg Mountain near Caledon, South Africa (34°12’S 19°32’E; Fig. 1). The area is characterized by Overberg Sandstone and Western Coastal Shale Band Vegetation Fynbos (Mucina *et al.* 2012) and hosts the entire *A. rugosa* population. Structurally, overstorey proteoid and asteraceous shrubs dominate the Klein Swartberg Mountain, creating an evergreen, fire-prone mosaic of open, mid-dense and closed vegetation coverage (Cowling *et al.* 1997; Mucina *et al.* 2012).

*Arthroleptella rugosa* habitat patches were identified based on previous records (Fig. 1), habitat type and thorough area searches. Searches were conducted in areas of recognized suitable habitat and consisted of two researchers walking approximately 50 m apart listening for *A. rugosa* calls (blue polygons; Fig. 1). *Arthroleptella rugosa* populations occur in small habitat patches sporadically dispersed along the main mountain ridge (green and red polygons; Fig. 1). The area of identified *A. rugosa* patches, and thus the actual spatial extent of their distribution, was determined to facilitate an accurate calculation of the total population size using estimated densities.

### Acoustic Surveys

Seven sites ranging from the eastern to the western side of the *A. rugosa* distribution were selected on the basis of logistical feasibility for acoustic surveying and were sampled from July through to September 2015 to coincide with the breeding season and associated peak calling times (green polygons; Fig. 1).

Each site was surveyed three times to correct for imperfect detections associated with fluctuating environmental conditions or survey time. For each survey, six Audio-Technica AT8004 Handheld Omni-directional Dynamic Microphones were connected to a DR-680 6-Track Portable Audio Recorder and set to record to six independent but synchronous tracks with a resolution of 24-bit and a recording frequency of 48 kHz. Microphones were attached to 1 m wooden dowels that were inserted into plastic tubing arranged in an array positioned in the centre of the habitat patch of calling *A. rugosa* males. The plastic tubing remained at each site between surveys to keep microphone positions constant throughout the survey period. Straight-line distances between all microphone pairs were measured to the nearest centimetre and the GPS locations (Garmin GPSMap64) of each microphone were recorded. Once recording commenced, the immediate survey area was vacated, and *A. rugosa* males were recorded for 40 minutes. Surveys were conducted exclusively in good weather conditions (no rain; minimal wind) to maximize individual call detectability (see Measey *et al.* (2017) for an explanation and more details on methodology).

### Data Pre-Processing

Raw acoustic recordings were processed to identify *A. rugosa* detections using the open source software for Passive Acoustic Monitoring, PAMGuard (Gillespie *et al.* 2009). The PAMGuard *Click Detector* was configured to detect the *A. rugosa* advertisement call using characteristics identified in Turner & Channing (2008). For each detection, the start time, relative amplitude and identification number of the microphone that made the detection were recorded. The start time for each call was recorded with an accuracy of 2.083×10^−5^ seconds.

### Acoustic Spatially Explicit Capture-Recapture (aSCR)

The capture histories (call start time, relative amplitude and detection information) obtained during pre-processing were used to determine the *A. rugosa* calling male density (frogs per hectare) using acoustic spatially-explicit capture-recapture (aSCR), a novel methodology outlined in Stevenson *et al.* (2015) and further applied in Measey *et al.* (2017). In processing the PAMGuard detection outputs, the first 5 minutes of each recording were omitted from the data to account for disturbance to the survey area caused by the presence of researchers. The detections (and non-detections) of calls across the array, and associated amplitudes and start times for each detection, facilitated the approximation of frog call locations. Detection probability functions were estimated using call and microphone locations (collected in the field, but corrected according to the distances between microphone pairs for precision; see Measey *et al.* 2017), and used to determine the probability that a call emitted from any location in the survey area was detected by at least one microphone. From this, the proportion of calls detected across the survey area and the estimated area in which these calls were made over the survey period (Effective Survey Area; ESA) were calculated. The call density was estimated by dividing the total number of calls by the ESA and the survey length.

The use of aSCR requires an understanding of the target species vocal behaviors (Marques *et al.* 2013). Specifically, accurate vocalization call rates estimates are necessary to precisely estimate calling animal density. During each acoustic survey, individual males were located and recorded for 20 minutes using a directional hand-held recorder (Olympus LS-10) to attain samples of *A. rugosa* advertisement calling (n = 22). Recordings of individuals were manually annotated in the open source software Audacity (see http://audacityteam.org/) to determine the number of *A. rugosa* advertisement calls made per minute.

### Incorporation of Population Boundaries

In previous aSCR studies, each survey area has been defined by a buffer around the microphone array (Stevenson *et al.* 2015; Measey *et al.* 2017; e.g. Fig. 2a). This assumes that calling males are present throughout this extent. Knowing, however, that the distribution of *A. rugosa* males does not conform to this spatial arrangement, this study delineated the actual patch area, determined during area searches by delineating the area with the path function on a Garmin GPSMap64, as the survey area (Fig. 2b). This masking process improves the precision of call location estimates, and thus also the detection function and density estimates.

**Figure 2.**
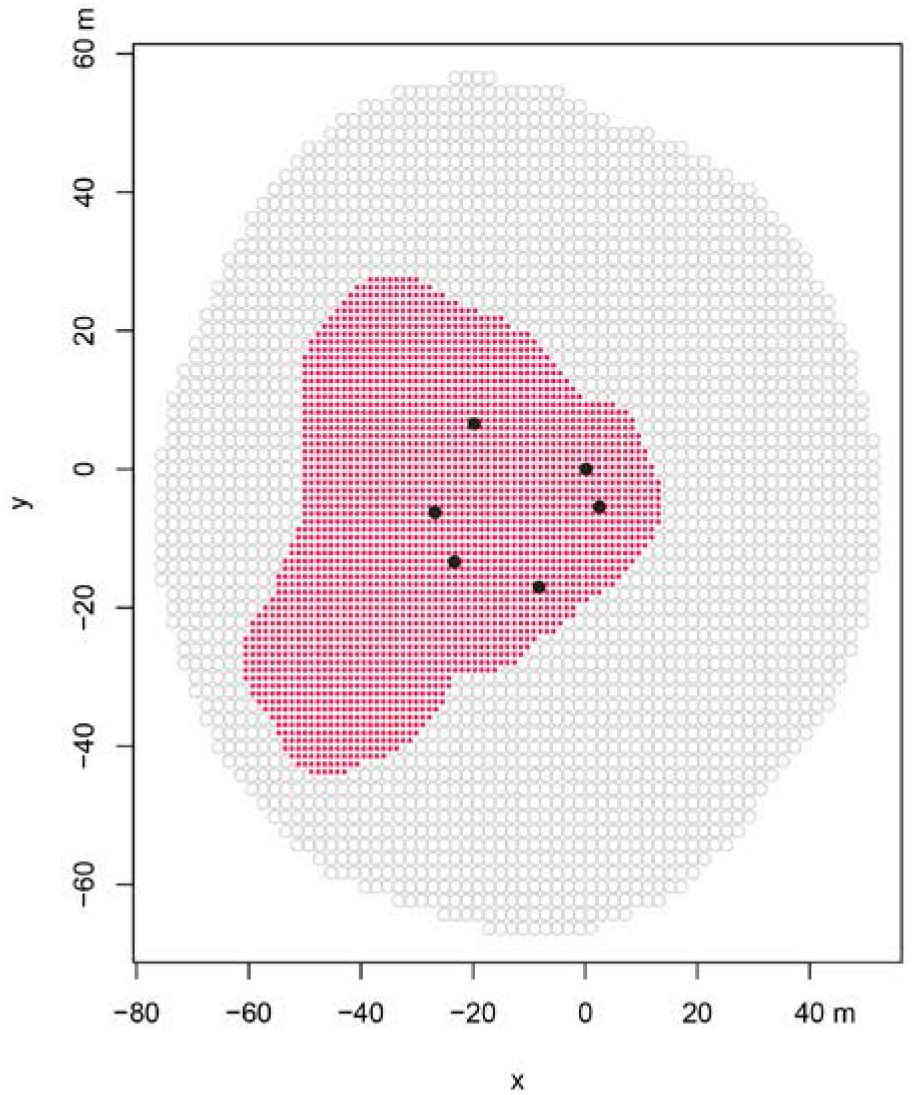
Graphical representation of the previously used 40 meter buffer of assumed frog presence (grey circles) and an example of the input used to delineate the extent based on suitable habitat (red circles). Black dots represent exact placement of microphones in the field, with x and y co-ordinates in meters.

#### Population Estimates

The R package ascr (Stevenson & Borchers 2015) was used to determine the detection function estimates and the call density. Modelling was computationally intensive, and so ten 2-minute subsamples were analyzed for each recording, giving a total of 20 minutes of analysis.

The density estimate for each surveyed patch was multiplied by the relevant patch area (in ha) to obtain site-specific population size estimates. The population sizes for the 11 unsurveyed habitat patches were determined by multiplying the mean density from the seven surveyed sites by the total area of the unsurveyed *A. rugosa* habitat (2.42 ha). The total *A. rugosa* adult population size across the Klein Swartberg Mountain was a summed estimate of population sizes from all surveyed and unsurveyed populations (multiplied by two to account for non-vocalizing females, assuming a sex ratio of 1:1). We acknowledge that bias in this estimation of the total population is inherent as we were not able to randomly sample *A. rugosa* habitat patches.

#### Quantifying uncertainty in density estimates

A parametric bootstrap included in the *aSCR* package was used to estimate the uncertainty of the density estimates calculated. Call rate data were included in the bootstrap simulation to account for the dependence between call locations and to attain the calling male densities with standard errors for each survey area. The bootstrap procedure was run for 500 – 1,000 iterations. To correct for inconsistent results between bootstrap-procedure repetitions, a secondary bootstrap was run concurrently for 500 iterations to quantify the *Monte Carlo Error* (MCE) associated with the density and standard error estimates generated (Stevenson, 2016). The number of bootstrap iterations for each analysis was increased until the Monte-Carlo error was below 0.05, where the MCEs for all parameters are not more than 5% over their standard error. This ensured that the number of bootstrapping iterations were sufficient to provide accurate parameter estimates.

The estimation of total population size was calculated as a sum of the estimated populations within surveyed areas and the estimated population within the unsurveyed area. The population estimate for each surveyed area was calculated using the average density and associated standard error for that site from aSCR estimates, whereas the *A. rugosa* populations within unsurveyed habitats were calculated using the reported average density of all sites surveyed, and the Standard Error of this total population was calculated from the difference of all density estimates, ignoring the error generated for each individual site.

### Habitat Assessments

Habitat assessments were carried out in each survey area to (i) identify the typical features of *A. rugosa* habitat and (ii) assess *Pinus pinaster* and *Hakea sericea* invasion. Ten 1 m^2^ quadrats were randomly placed throughout each survey area. For each quadrat, the percentage of ground cover (i.e. grasses and reeds) within the quadrat was estimated and the heights of at least three plants occurring within the quadrat were recorded. Where alien vegetation was present within a survey area, the number and height of alien plants were also recorded,. Additionally, the level of invasive vegetation (adult and these werejuvenile plants) present surrounding the survey area was categorized into low (<5 plants), medium (5-15 plants) and high (>15 plants).

### Statistical Analysis

Summary statistics were carried out to identify typical *A. rugosa* habitat characteristics. A linear model was performed in R (version 3.5.1) to assess the impact, if any, of invasive vegetation on male calling densities. Invasive vegetation features included in the analysis were: presence/absence of invasive species within the recording site, and the level (low/medium/high) of invasive vegetation adjacent to the recording site.

### Ethics Statement

This study was carried out within the permission of the provincial conservation authority, CapeNature (Permit Number: AAA007-00162-0056) and was approved by the Animal Ethics Committee of the University of Queensland (Permit Number: ANRFA/SBS/280/15/STELLENBOSCH). Fieldwork was carried out on privately owned land, with access granted by land owners. The species targeted in acoustic surveys was a Critically Endangered species of frog, *Arthroleptella rugosa*. The survey methods chosen for this project were selected specifically to minimize researcher disturbance to the *A. rugosa* habitat, and extra care was taken in the field to ensure no unnecessary disturbance occurred.

## RESULTS

Area searches for calling *A. rugosa* males yielded the discovery of 12 previously unknown *A. rugosa* populations. In total, calling individuals occupied 5.15 ha of land across the Klein Swartberg Mountain, representing 2.2% of the 230 ha suitable area. The acoustic surveys (n = 7 sites) covered over 50% (2.74 ha) of this area.

There was an average of 35,205 ± 4,505 detections of *A. rugosa* calls made during each 40 minute recording, of which a total of 16,962 ± 1,740 detections were included in the 2-minute subsamples (n = 10) used in the final analysis. The habitat patch area (in hectares) and the mean number of calling males per hectare (with their associated standard errors) for each site surveyed are displayed in Table 1.

**Table 1.**
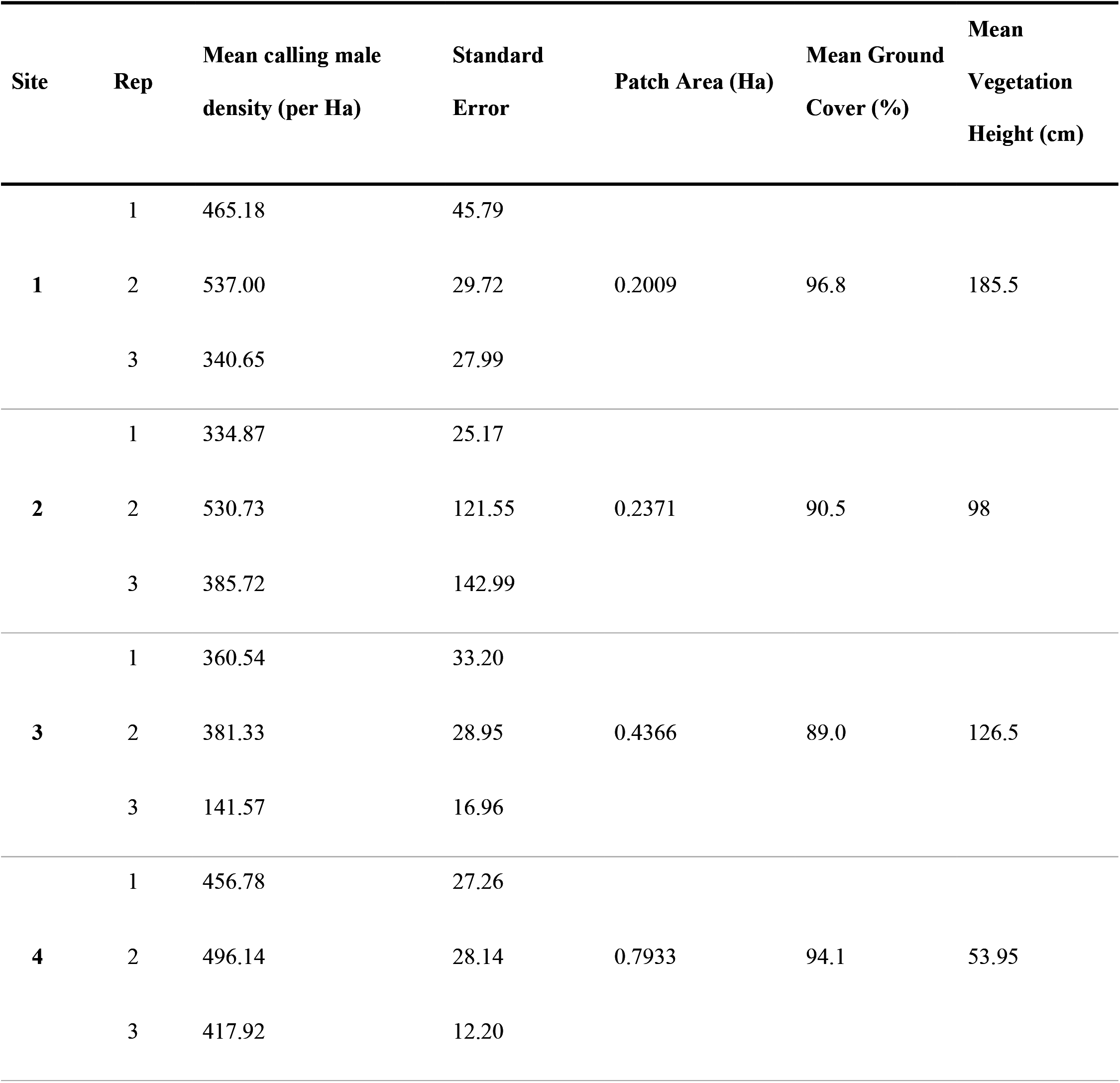
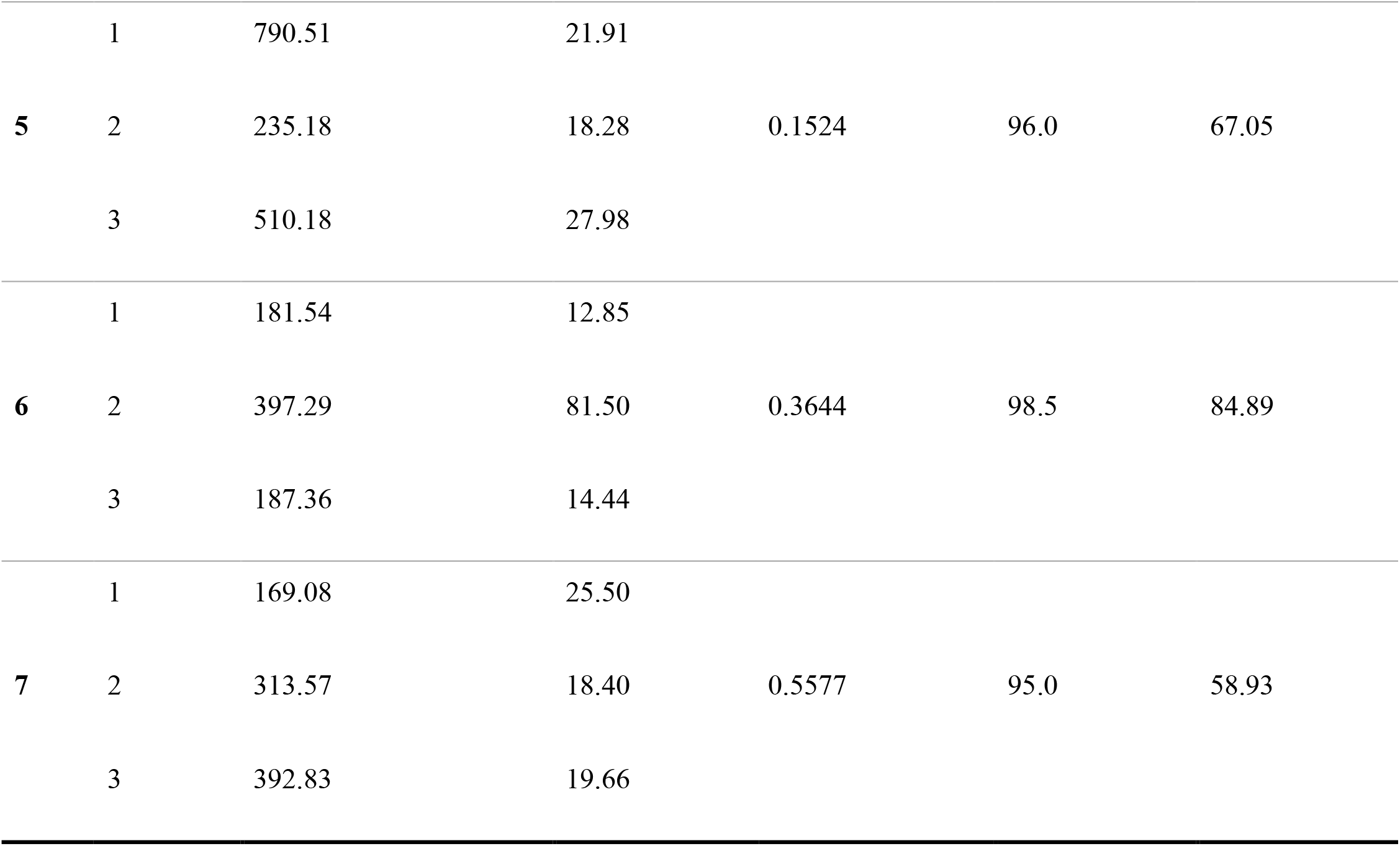
Results of vegetation assessments and the mean *Arthroleptella rugosa* calling male density per hectare and associated standard error (SE) for each replicate (n = 3) of all seven surveyed sites. The patch area (in Hectares), mean ground cover (%) and mean vegetation height (cm).

The mean advertisement call rate was 13.49 ± 0.97 calls per minute.

Across all survey sites, the total mean calling male density was 417 individuals per hectare (± 21.7 individuals per hectare), and the estimated male population size in the surveyed area (2.74-ha) was 1,053 individuals (± 194.1 with 95% CI). When considering both the survey area and the surrounding unsurveyed area (i.e., the total occupied area; 5.15-ha), the estimated total male population size was 2,060 individuals (± 132.2 with 95% CI). Therefore, assuming a sex ratio of 1:1, the total adult population size is estimated to exceed 4,000 individuals.

### Habitat Assessments

The survey sites representing *A. rugosa* habitat were characterized by an average native vegetation height of 97.2 cm (± 8.2 cm; 95% CI) and 94.3% (± 1.3%; 95% CI) ground cover. All seven survey sites were adjacent to streams that dispersed water throughout the seepage area, maintaining a moist environment. Sixteen *P. pinaster* saplings, with an average height of 46.8 cm (± 6.8 cm; 95% CI), were found within survey site 3 (western-most surveyed site within the eastern cluster of survey sites). No other surveyed areas were invaded by alien vegetation, although much of the area surrounding surveyed sites within the western portion of the species’ distribution (sites 1-3) was invaded by adult and juvenile *P. pinaster* and *H. sericea*. Significantly higher male calling densities were found at sites adjacent to low or no invasive vegetation, and no invasive species present in the surrounding habitat (p < 0.0001, F_3,211_=23.82, R^2^ = 0.2424; Fig. 3).

**Figure 3.**
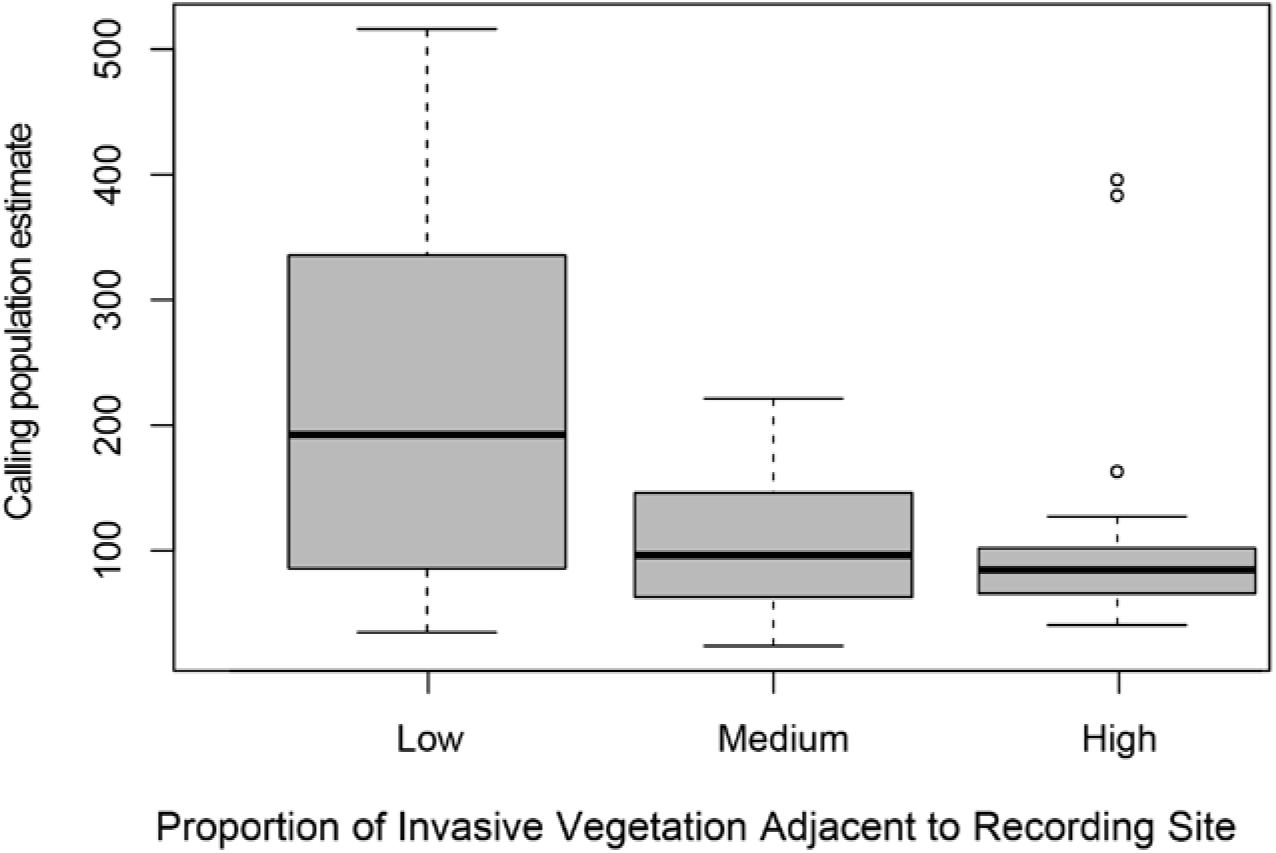
The difference in calling densities of *Arthroleptella rugosa* from calling sites with different quantities of invasive pine and hakea in adjacent habitat in the Klein Swartberg, South Africa.

## DISCUSSION

The *A. rugosa* total adult male population size was estimated to be 2,060 individuals (± 132.2 with 95% CI) using the newly developed acoustic spatially explicit capture-recapture (aSCR) technique. This estimate is more than five times higher than that formerly predicted by Turner and Channing (2008), where aural calling surveys projected the population to be 400 adults. At a sex-ratio of 1:1, the total adult population is estimated to exceed 4,000 individuals. This practical application of aSCR demonstrates how baseline population data can be determined for an entire species, providing population estimates that can be used in conservation assessments. Moreover, the methodology allows for repeated surveys with statistically robust estimates for monitoring purposes, where individual populations of the species can be targeted for repeated recordings (see Measey *et al.* 2017). These could be used to track the real impact of threats to species, such as alien plants and fire.

*Arthroleptella rugosa* populations are distributed within 2.2% of habitat patches across 230 ha of suitable habitat on the Klein Swartberg Mountain. Survey areas were, on average, characterized by high ground coverage and vegetation shorter than 1 m. Considering a large portion of the *A. rugosa* habitat was affected by fires in January 2012 (Measey *et al.* 2019), and fires have been associated with devastating adult frog population declines (Channing, 2004), the estimates from this study suggest that the *A. rugosa* population has started to recover, even in dense alien-invaded habitat where the destruction from fires would be most intense. However, it is likely though that this apparent increase in population size was augmented by differences in the sampling techniques used and the additional populations discovered. The total population estimate from the present study was estimated by summing the population sizes for each surveyed and unsurveyed habitat patch, which covered the total patch area of 5.15 ha, even though the patches were not randomly sampled. In contrast, Turner and Channing (2008) conducted aural calling surveys within the six populations (2.78 ha) known at the time. The discovery, and subsequent inclusion, of the additional 2.37 ha of *A. rugosa* habitat in the present study would likely lead to higher population estimates. Alternatively, the currently implemented *P. pinaster* and *H. sericea* control across much of the Klein Swartberg Mountain could be adequately maintaining waterways, streams and seepage areas, sustaining suitable habitat for *A. rugosa* and facilitating steady population increases.

Although the total population appears to have increased from the previous estimation, the entire population of *A. rugosa* appears to be fragmented. A large *P. pinaster* stand the eastern and western clusters located in the middle of surveyed sites the Klein Swartberg Mountain ridge (see Fig. 1; largest yellow polygon) could be acting as a barrier, physically and hydrologically separating the western populations from the remaining central and eastern populations. This would restrict the successful movement between and re-colonization of suitable habitat patches, affecting the dynamics of the meta-population and threatening *A. rugosa* persistence (Marsh & Trenham 2001). The hydrological distance between populations has been described as one of the most confounding factors affecting dispersal in small-sized mountain-dwelling anurans, as individuals tend to travel further along streams than through terrestrial spaces typically interspersed with unsuitable habitat (Measey *et al.* 2007). For direct-developing species like *A. rugosa*, where populations can occur continuously within habitats, extend past the ‘boundaries’ of suitable habitat areas and even occur throughout fragmented habitats (Measey *et al.* 2007), maintaining passive and active dispersal is integral to their conservation. As such, recognizing the importance of streams in maintaining connectivity across the *A. rugosa* distribution is essential to long-term population persistence.

Total population size and site occupancy of anurans can be highly variable, and so consistent, long-term monitoring of the *A. rugosa* population is necessary to assess population fluctuations and trends in relation to various conservation actions, environmental conditions and other influential factors (Breven 1990). For *A. rugosa*, the effects of emerging invasive pines and hakea on *A. rugosa* populations might not be immediately apparent, however, the direct effects of fire would be. This time lag may lead to difficulties in determining and targeting conservation efforts to mitigate all threatening processes affecting *A. rugosa*, as initially some threats may appear more severe than others (De Solla *et al.* 2006). Where possible, population survey information should be derived from long-term monitoring data.

The currently implemented management actions and higher than anticipated *A. rugosa* population size are evidence of the value of private reserve areas to amphibian conservation. The Klein Swartberg Conservancy is a designated 14,857 ha area where both landowners and CapeNature officials have successfully implemented conservation actions directed at preserving *A. rugosa* populations and habitats. This conservancy provides the much-needed protection and restoration of not only *A. rugosa* habitat areas, but also the water systems that are critically endangered in the context of the broader landscape and the many other species that occur in this area. Thus, this conservancy is contributing directly to the conservation of a critically endangered species, and to the maintenance of waterways in the area. Monitoring of *A. rugosa* now requires the adoption and application of the aSCR method by CapeNature.

While the aSCR method accounts for distribution differences across the ESA, as estimates are averaged and extrapolated across all sites, this study assumes that the frogs are uniformly distributed across the occupied landscape. Population densities across the *A. rugosa* distribution, and thus also local population sizes, can vary significantly, as the landscape, altitude and habitat characteristics are not homogenous across the Klein Swartberg Mountain. This is evidenced within this study by the stark differences observed between the densities obtained for each of the surveyed sites (Table 1). Within survey areas, individuals may be unevenly distributed as they may associate with certain habitat features. Therefore, at both the survey area and full distribution scales, this assumption is likely to be violated. The present study attempted to address this through the novel incorporation of population bounds, improving the precision of population estimates from each survey area. Furthermore, biases to the total population estimate would be mitigated through the inclusion of multiple survey areas to attain a representative sample of population densities across the *A. rugosa* distribution.

This method would be easily adoptable across various habitat types and species, as its implementation in the field does not require explicit user-training or expert species knowledge. It also has relatively low survey effort and can be adapted to suit a variety of spatial scales and to simultaneously monitor multiple species. Thus, aSCR presents the opportunity to implement monitoring to assess conservation efforts in both on and off reserve areas that have previously gone unchecked. Monitoring populations in these areas can identify the outcomes of implemented conservation actions, and thus audit their success. In this study, aSCR was used to rigorously estimate *A. rugosa* population densities, setting the precedent of using this technique on small, cryptic, vocally distinct species and providing the basis for commencing adequate monitoring schemes for many data deficient amphibian species. While this method is an effective conservation tool that can be used to maximize the efficiency of management actions, there are still many opportunities for further development. Advances in the incorporation of animal movement or individual animal recognition would significantly improve the precision of estimates and allow this method to be applied to any vocally distinct species. Furthermore, transitioning to a wireless, automated system would facilitate an easier collection of data without the presence of an observer. As aSCR methodologies are refined, this method will become an essential conservation tool used for monitoring acoustically active species.

## Acknowledgements

We thank Carel le Roux and Michael Swart for access to their properties. DS & JM thank the DSI-NRF Centre of Excellence for Invasion Biology for support. AT & JM thank Atherton de Villiers for his help in conducting regular monitoring of *Arthroleptella rugosa* on the Klein Swartberg Mountain and the US Fish and Wildlife Service for Grant F11AP00335. Equipment used in this study was funded by the National Geographic Society/Waitt Grants Program (No. W184-11) to JM.

## Supplemental Material

Original acoustic audio files: South African Environmental Observation Network (SAEON) Data Repository http://dx.doi.org/10.15493/SAEON.METACAT.100000XX (Measey *et al.* 2018). To be added on acceptance of ms.

